# Striatin is required for hearing and affects inner hair cells and ribbon synapses

**DOI:** 10.1101/2020.03.11.987396

**Authors:** Prathamesh Thangaraj Nadar Ponniah, Shahar Taiber, Michal Caspi, Tal Koffler-Brill, Amiel A. Dror, Moran Rubinstein, Richard A. Lang, Karen B. Avraham, Rina Rosin-Arbesfeld

## Abstract

Striatin, a subunit of the serine/threonine phosphatase PP2A, is a core member of the conserved striatin-interacting phosphatase and kinase (STRIPAK) complexes. The protein is expressed in the cell junctions between epithelial cells, which play a role in maintaining cell-cell junctional integrity. Since adhesion is crucial for the function of the mammalian inner ear, we examined the localization and function of striatin in the mouse cochlea. Our results show that in neonatal mice, striatin is specifically expressed in the cell-cell junctions of the inner hair cells, the receptor cells in the mammalian cochlea. Auditory brainstem response measurements of striatin-deficient mice indicated a progressive, high-frequency hearing loss, suggesting that striatin is essential for normal hearing. Moreover, scanning electron micrographs of the organ of Corti revealed a moderate degeneration of the outer hair cells in the middle and basal regions, concordant with the high-frequency hearing loss. Importantly, striatin-deficient mice show aberrant ribbon synapse maturation that may lead to the observed auditory impairment. Together, these results suggest a novel function for striatin in the mammalian auditory system.

## Introduction

The striatin-interacting phosphatase and kinase (STRIPAK) complex is a multimolecular protein complex involved in numerous biological functions and has been implicated in a number of human diseases [1–3]. STRIPAK complexes regulate phosphorylation of diverse proteins and interact with conserved signaling pathways [2]. The mammalian striatin family is ubiquitously expressed and consists of striatin (STRN), SG2NA (STRN3), and Zinedin (STRN4). All these proteins contain multiple WD40 repeats, as well as a Ca^2+^-calmodulin-binding domain, a caveolin-binding motif, and a coiled-coil structure that are essential for their function [4, 5]. We have recently shown that striatin is expressed at cell-cell junctions and is required for cell-to-cell adhesion [6].

Within the cochlea, the mammalian inner ear contains anatomically and functionally distinct inner hair cells (IHCs) and outer hair cells (OHCs), which convert mechanical stimuli into electrical signals [7]. The different types of cell-cell junctions and proteins within the cochlea maintain the well-organized structure and function of the inner ear and enable hearing.

Here, we examined the localization and function of striatin in the mouse inner ear in order to evaluate its role in the auditory system. Our results show that striatin is specifically expressed in cell junctions between the IHCs of the organ of Corti, and that striatin knockdown leads to progressive hearing loss. Although loss of striatin did not lead to changes in the junctional integrity of the hair cells, or affect the endocochlear potential (EP), partial degradation of the OHCs was observed, particularly in the basal region of the organ of Corti. This observation is consistent with our results showing that striatin-deficient mice exhibit high-frequency hearing loss. Normal hearing requires proper positioning of the ribbon synapses that undergo complex organizational processes during the maturation of inner hair cells [8–11]. Our results show that striatin-deficient mice have a uniform distribution of ribbons that lack the spatial gradient seen in wild-type IHC. Interestingly, a similar aberrant maturation of the synapse is seen in striatin binding protein adenomatous polyposis coli (APC) mutant mice [12]. Taken together, our results present striatin as a novel multifunctional protein that is essential for mammalian hearing.

## Results and Discussion

The STRIPAK complex, which includes striatin, a subunit of the PP2A enzyme, is associated with numerous biological roles ranging from cell signaling to developmental processes [2]. To study the biological function of striatin, a knockout mouse was generated.

### Striatin is expressed in the apical surface of the inner hair cells

Depending on cell type and condition, mammalian striatin localizes to diverse subcellular compartments such as the Golgi [13], endoplasmic reticulum, plasma membrane, mitochondria [14, 15], and cellular junctions [6]. As cell-cell adhesion is essential for auditory function, and since striatin had low diffuse expression in the cochlear sensory epithelium in the early postnatal stages, with a higher expression detected in inner and outer hair cells in the maturing cochlea [16–18] (umgear.org), we were interested in examining the expression pattern and role of striatin in the auditory system. To this end, we designed and constructed a striatin-deficient mouse (Fig EV1). Cochlea of P0 mice were dissected and total protein was extracted for western blot analysis. The specific striatin knockdown was verified by western blot analysis of wild type (WT), *Strn*^*+/-*^ and *Strn*^*-/-*^ littermates (Fig EV2A). As expected, there was no immunostaining of striatin in the *Strn*^*-/-*^ mice. We further examined the expression and localization of striatin in the inner ear; a schematic illustration of the cochlea, including the localization of striatin, is shown in Fig 1A. Results shows that striatin is indeed expressed in the cochlea at the protein level (Fig 1B). The localization of striatin in the organ of Corti was evaluated by immunofluorescence assays. Here, striatin was specifically detected in the cell-cell junctions of the IHCs (Fig 1C, D).

**Figure 1.**
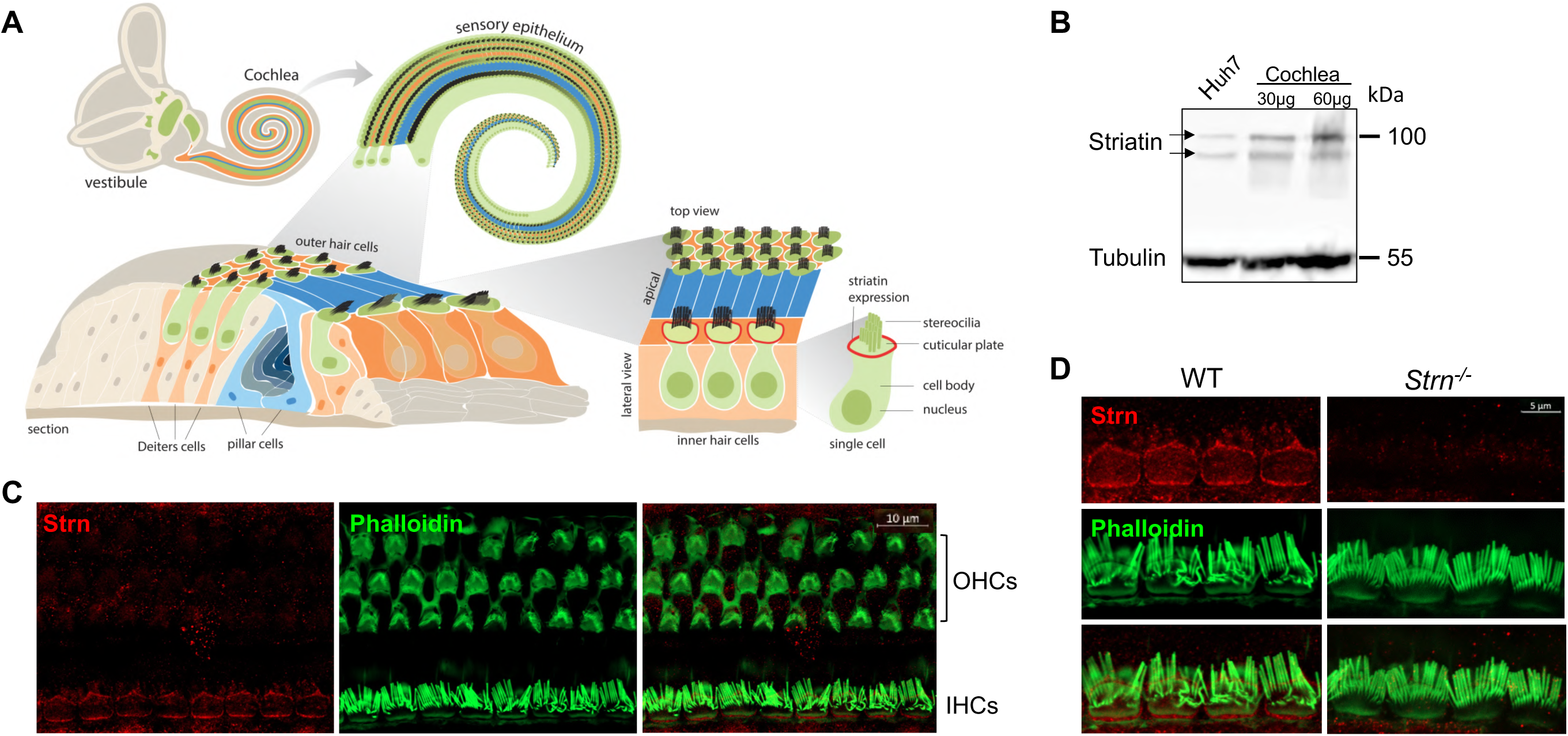
Striatin is expressed in cell-cell junctions of the mouse inner hair cells. A Schematic representation of the mouse inner ear. B Striatin is expressed in the cochlea. Cochlea and Huh7 (as a positive control) cell lysates were analyzed by SDS-PAGE and immunoblotted using the indicated antibodies. Tubulin served as a loading control. C Striatin is expressed in the inner hair cells (IHC). Immunofluorescence analysis of striatin expression in P30 whole-mount mouse inner ears stained with striatin, and phalloidin to visualize filamentous actin. D Striatin is localized to the sites of cell-cell junctions in IHCs. Immunofluorescence analysis performed on WT and *Strn*^*-/-*^ litter mates show that striatin is localized to the sites of cell-cell junctions between the IHCs.

### Striatin knockout mice exhibit progressive hearing loss

To further understand the function of striatin in the auditory system, we used the Auditory Brainstem Response (ABR) recordings. ABR was performed on P20, P30, P40, and P60 WT, *Strn*^*+/-*^, and *Strn*^−/-^ littermates using a sound stimulus with varying frequencies (6–35 kHz) and intensities (10–90 dB). The results showed that the hearing threshold for *Strn*^*+/-*^ mice was similar to that of the WT mice at P20, and that there was a modest shift in hearing thresholds for certain frequencies in the *Strn*^−/-^ mice compared to the WT mice. However, at P30, there was a substantial increase in the hearing thresholds of both the homozygous and heterozygous striatin mice, compared to the WT mice. At P40 and P60, both *Strn*^*+/-*^ and *Strn*^*-/-*^ mice showed severe hearing loss, indicating that striatin deficiency leads to progressive hearing impairment. Representative ABR traces in response to 30 kHz sound stimuli is presented in Fig EV3. The striatin mutant mice had compromised hearing at frequencies above 12 kHz, implying that the mid and basal regions of the organ of Corti could be affected.

### Striatin mutants show moderate outer hair cell degeneration

To better understand the observed hearing loss, we examined hair cell degradation using scanning-electron microscopy (SEM). The results show moderate OHC degeneration in *Strn*^*+/-*^ and *Strn*^*-/-*^ mice as compared to WT mice (Fig 3A). Quantification of the hair cells revealed 21.6% and 9.9% loss of OHCs in the basal and mid regions of the cochlea, respectively, for *Strn*^*-/-*^ mice, and 13% and 6.8% loss of OHCs in the basal and mid regions, respectively, for *Strn*^*+/-*^ mice (Fig 3B). Although OHC loss was not substantial, this might contribute to the higher frequency hearing loss of the *Strn*^*-/-*^ mice. However, these results imply that hair cell degradation is not the only cause of the severe hearing loss we observed. OHC loss could also be secondary to initial damage to other components of the auditory system.

**Figure 2.**
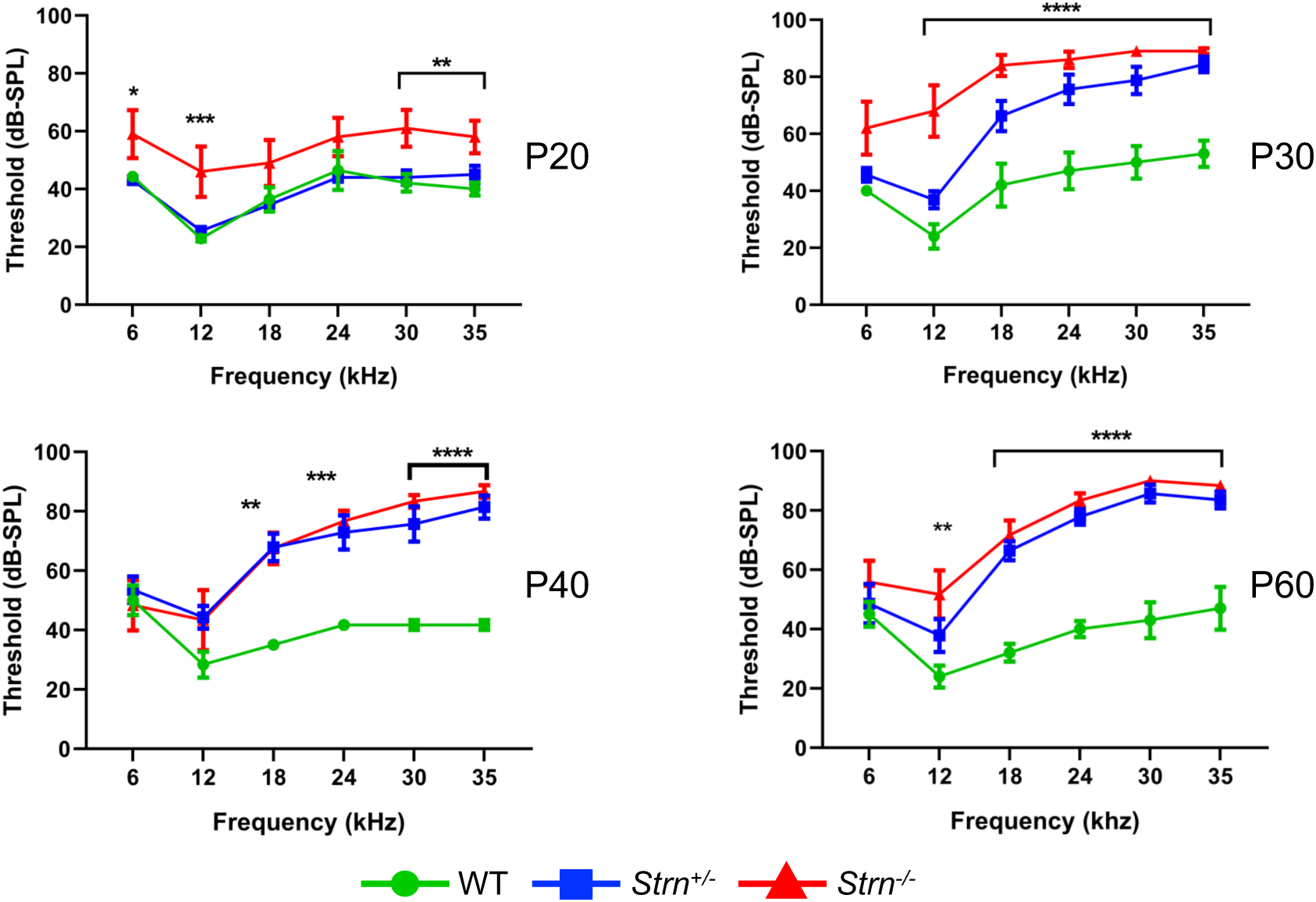
Auditory brainstem response (ABR) reveals progressive hearing loss in *Strn*^*-/-*^ mice at higher frequencies. ABR threshold means are shown for *Strn*^*-/-*^ mice and littermate control mice, tested at the indicated ages. P20: n = 10 *Strn*^*+/-*^, 5 *Strn*^*-/-*^, and 7 WT, P30: n = 8 *Strn*^*+/-*^, 5 *Strn*^*-/-*^, and 5 WT, P40: n = 7 *Strn*^*+/-*^, 6 *Strn*^*-/-*^, and 3 WT, and P60: n = 7 *Strn*^*+/-*^, 6 *Strn*^*-/-*^, and 5 WT. Data shown as mean ± SEM. Statistical tests were two-way ANOVA with Holm-Sidak multiple comparison correction. *P < 0.1 **P < 0.01 ***P < 0.001 ****P < 0.0001

**Figure 3.**
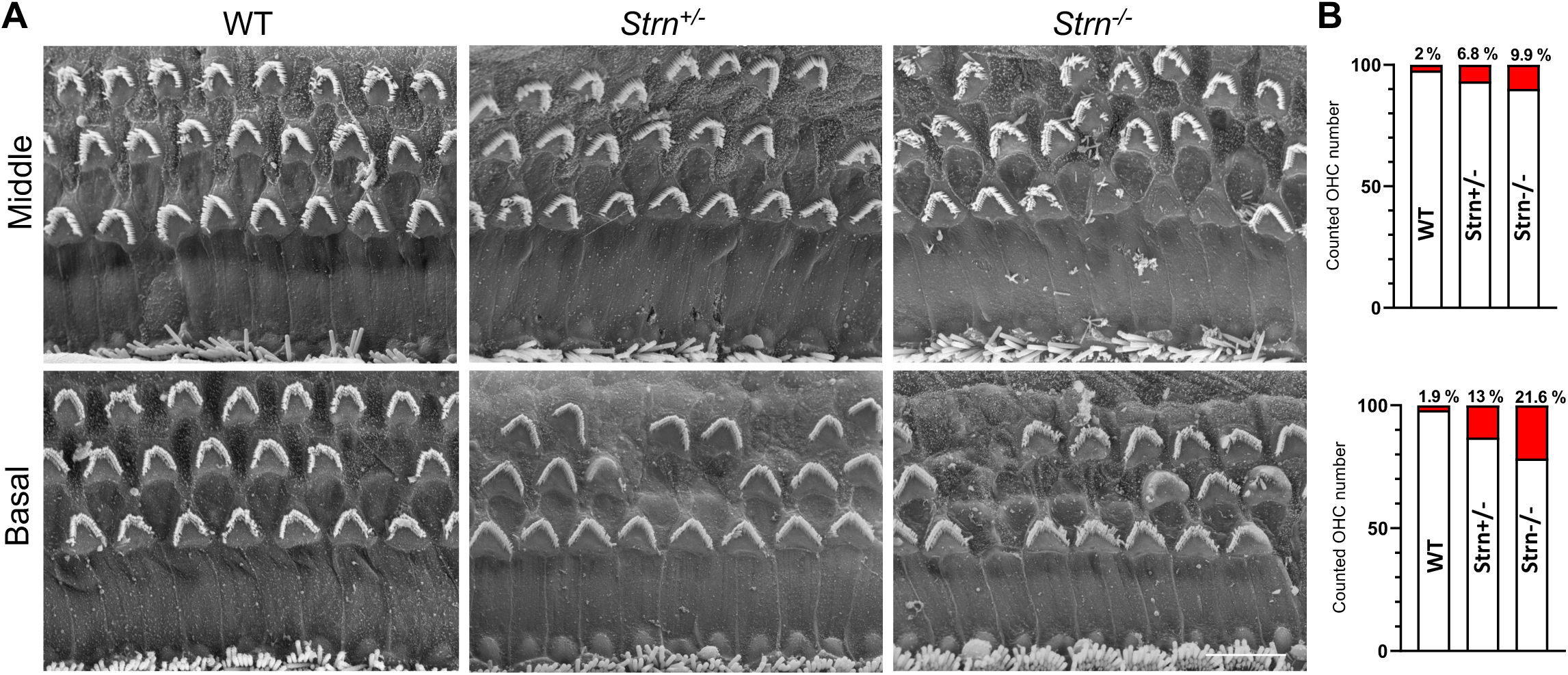
Scanning electron micrograph and quantification of hair cells show moderate outer hair cell (OHC) degeneration in *Strn*^*-/-*^ mice. The hair cells were counted in 200 μm stretches of middle and basal coil regions at P60. A Representative electron micrograph reveals moderate hair cell degeneration of OHC in *Strn*^*+/-*^ and *Strn*^*-/-*^ as compared to WT in the middle and basal region of the organ of Corti. B The graphs show numbers of cell loss/survival for middle and basal regions. Middle coil region (upper panel): 2/87 (Control, n = 3), 7/103 (*Strn*^*+/-*^, n = 4), 28/284 (*Strn*^*-/-*^, n = 12), and basal coil region (lower panel): 4/206 (Control, n = 8), 43/274 (*Strn*^*+/-*^, n = 11), 32/148 (*Strn*^*-/-*^, n = 7). Cell loss is marked in red.

### Cell-cell junctional integrity is retained in the striatin knockout mice

Cell-cell junctions in the cochlea are crucial for maintaining the correct structure and function of the organ of Corti and complete or partial loss of function of cell junction proteins often results in impaired hearing [19–22]. Striatin was shown to maintain junctional integrity in cultured mammalian cells [6] which accords with our finding of striatin in IHC cell junctions (Fig 1). Immunostaining was used to monitor the expression pattern and subcellular localization of the tight junctional (TJ) protein ZO-1 and the adherence junction protein E-cadherin in the hair cell junctions of striatin knockout mice. Interestingly, the expression pattern of both ZO-1 and E-cadherin in the knockout mice was similar to that of WT mice, indicating that the junctional integrity in the organ of Corti is maintained in striatin-deficient mice (Fig 4A and 4B). One possible explanation may be the presence of a redundant protein compensating for striatin loss. Striatin 4 shares over 50% protein sequence homology with striatin and also binds the catalytic subunit of PP2A, indicating that the proteins may have redundant cellular functions, as seen in various systems [1, 23].

**Figure 4.**
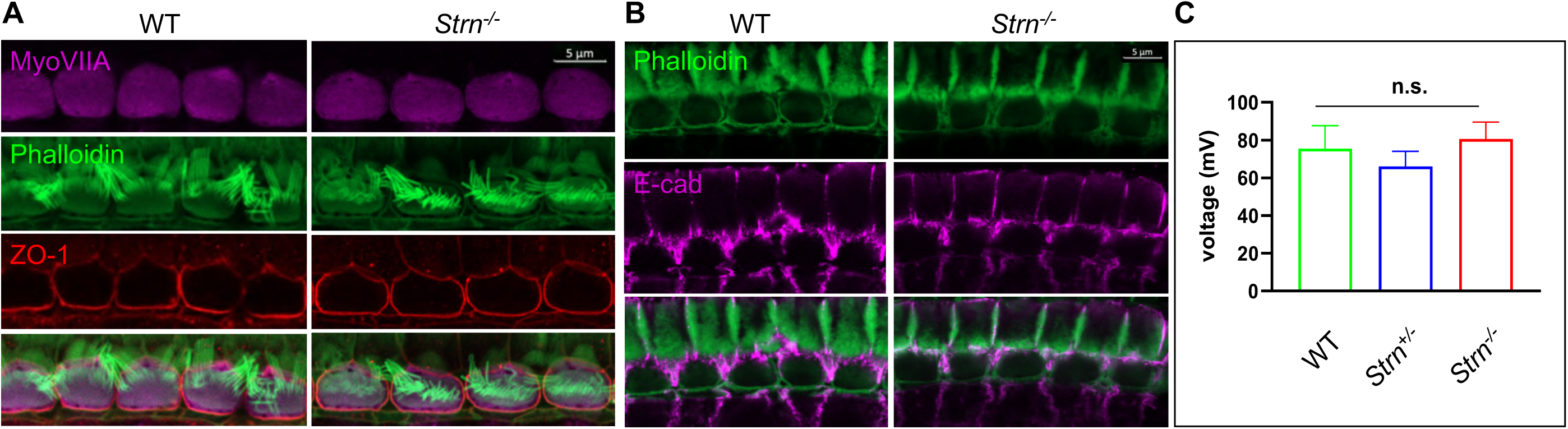
Junctional integrity in the organ of Corti is intact in *Strn*^*-/-*^ mutants. A, B Representative P30 whole-mount mouse inner ear stained with myosin VIIa (purple), ZO1 antibody (red) and phalloidin (green) (A), or E cadherin (purple) and phalloidin (green) B. The experiments were repeated using 4-5 mice of each genotype. C Recordings of endocochlear potential (EP) of endolymph are shown for *Strn*^*-/-*^ and littermate control mice, tested in 3-month old mice (*Strn*^*-/-*^ (n = 2), *Strn*^*+/-*^ (n = 7) and WT (n = 3). Statistical test was one-way ANOVA with Holm-Sidak correction for multiple comparisons.

The stria vascularis produces the endolymph extracellular fluid [24] and generates an endocochlear potential (EP) that is essential for normal auditory function [25, 26]. The TJs in the organ of Corti are required to form the cation-junctional barrier between perilymph and endolymph and to maintain the EP [27, 28]. The presence of striatin at the junctions of the IHCs led us to examine whether striatin knockout mice are capable of maintaining the EP. No significant changes were detected in the EP of striatin-deficient mice (Fig 4C). This finding is in accordance with our results showing that the junctional integrity between neighboring IHCs is not compromised by striatin loss.

### Striatin knockout mice display an aberrant ribbon gradient

Striatin is a subunit of the serine/threonine phosphatase PP2A [5], and reduced PP2A activity impairs synaptic function [29]. Interestingly, lack of another member of the STRIPAK complex, striatin interacting protein 2 (Strip2), leads to a decrease in neural response amplitudes. Since the synaptic ribbons of IHCs express Ctbp2 [30, 31], we examined the expression pattern of Ctbp2 in P17 and P35 mice, time points at which striatin-deficient mice exhibit hearing loss. Z-stacks of at least eight continuous IHCs were imaged in WT and striatin null mice. Myosin VIIa was used to stain the hair cell body. The 3D structures were analyzed using Imaris software, followed by quantification of the number and localization of the Ctbp2 puncta with respect to the modiolar and pillar faces of the IHC. Fig 5 shows a representative 3D side view of the IHCs that were used to determine the puncta number. Although the total number of Ctbp2 puncta at the synaptic poles of the *Strn*^*-/-*^ mice at two time points, P17 and P35, was not significantly different from that of WT mice (Fig 5D), the Ctbp2 puncta in *Strn*^*-/-*^ mice showed a uniform distribution and lack the ribbon localization gradient towards the modiolar side of the IHC, as seen in WT mice that show a spatial gradient towards the modiolar face of IHC (Fig 5A, C). This finding suggests aberrant maturation of the ribbon structure in striatin-deficient mice, which subsequently could affect auditory function. Interestingly, in adenomatous polyposis coli (APC), another striatin-interacting protein, deficient mice, [32, 33], ribbon synapses were shown to lack a size gradient towards the modiolar face of the IHCs [12]. APC conditional knock-out mice show impaired auditory function, which probably results from aberrant afferent synapse ribbon size gradients. Similarly to the *Strn*^*-/-*^ mice, the IHCs of the APC knock-out mice show wild-type ribbon numbers but lack the normal ribbon size gradient [12]. As ribbon synapses mediate transmission between IHCs and spiral ganglion neurons (SGNs), our results lead us to propose that striatin, a component of the STRIPAK complex that interacts with the tumor supressor protein APC, regulates interneuronal synapses. This finding is supported by the observation that the *Drosophila* striatin homolog, CKA, facilitates axonal transport of dense core vesicles and autophagosomes in a PP2A-dependent manner [34]. Moreover, as striatin functions as a PP2A subunit, it is also plausible to speculate that lack of striatin will disrupt the phosphorylation patterns that are essential for precise auditory function [35–37].

**Figure 5.**
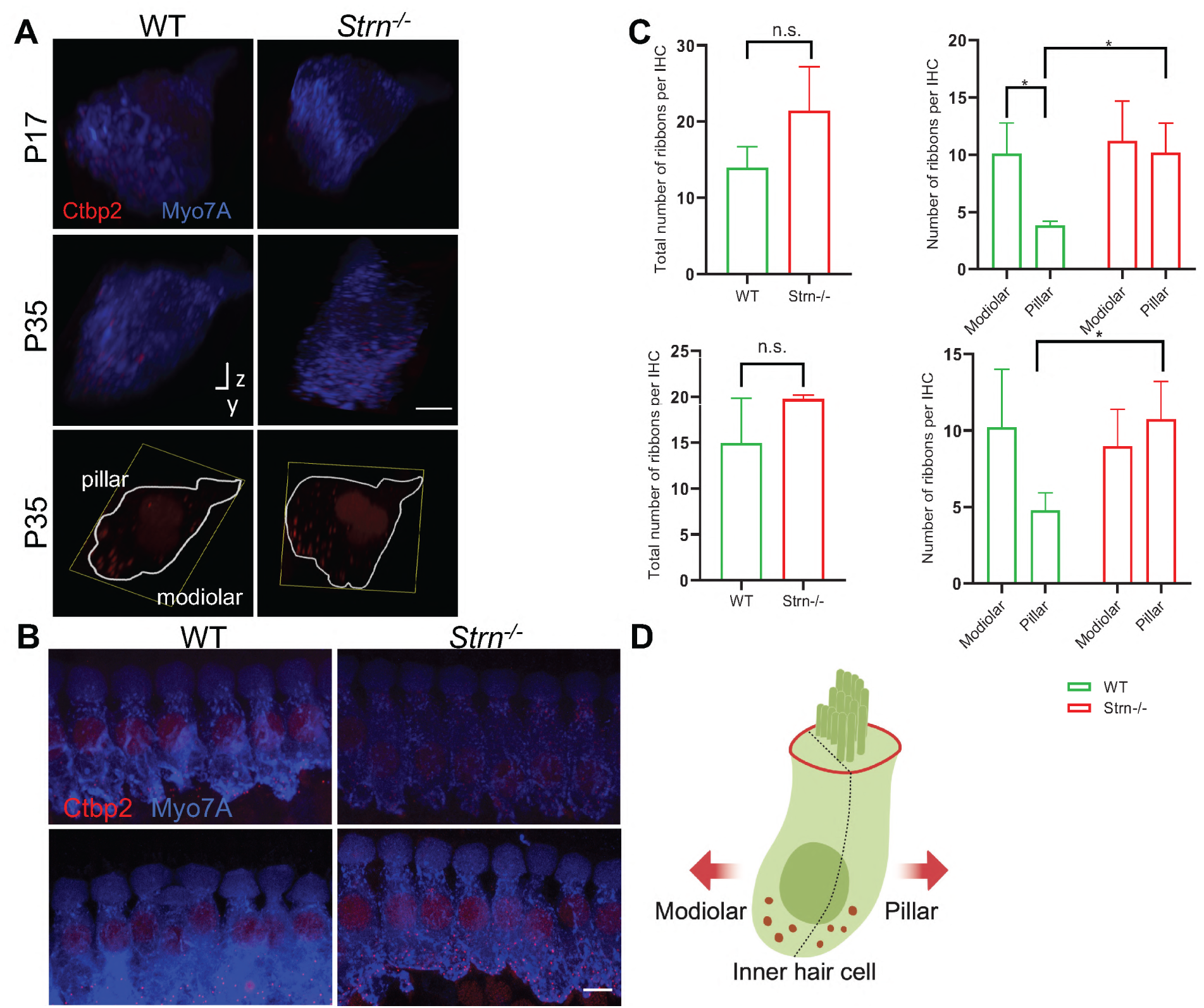
*Strn*^*-/-*^ mice lack a correct ribbon size gradient. Confocal micrographs of ribbons at the synaptic poles of individual IHCs (Ctbp2, red, and myosin VIIa, blue) in z-stacks, oriented on the y–z-axis. A Representative side view of at least 8 contiguous IHC in WT and *Strn*^*-/-*^ P17 and P35 mice. Bottom panels: representative side view of at least two IHCs stained at P35 with Ctbp2 (outlined in white based on myosin VIIa staining). B Representative maximum projection of myosin VIIa (blue) and Ctbp2 (red) in inner hair cells, with the latter staining synaptic poles of IHCs for WT and *Strn*^*-/-*^ P17 mice. C Total ribbon number and ribbon gradient along the modiolar and pillar regions in *Strn*^*-/-*^ mice and WT littermates. Imaris software was used to analyse the 3D z-stacks, 4-11 IHCs from each cochlea were imaged for quantitative ribbon analysis. Statistical analysis was performed by 2-way ANOVA and Holm-Sidak multiple comparisons test (n = 4). P17, upper panel; 035 lower panel. D Schematic diagram of an inner hair cell demonstrating plane of orientation of pillar vs modiolar region.

In this context, another STRIPAK component, Striatin interacting protein 2 (Strip2) [38, 39], which is expressed in OHC and IHC [16, 31, 40], is known to be important for neural response amplitudes [31]. In conclusion, our results provide the first evidence that striatin, a member of the conserved STRIPAK complex, functions in the auditory system. Although we hypothesize that the role of striatin may be related to synaptic transmission and cell-cell junctions, other roles are also possible, making this finding an exciting opportunity for further investigation and validation.

## Materials and Methods

### Strn-knockout mice: establishment and genotyping

The ES cell line EPD0082_3_E07 carrying the Strn^tm1a(KOMP)Wtsi^ allele was injected into embryos, which were transplanted into recipient C57BL/6 female mice. All animal procedures were approved by the Animal Care and Use Committee (IACUC) at Tel Aviv University (01-18-085). Genotyping was performed from tail samples by PCR, using a set of primers that flank the *Striatin* gene: F-5′TTCCTTTGAGAAAACACAGTCCCAG-3′, R-5′-ACACACTCCACTGAACAAAGTCAAGC-3′, to give a 1257bp product in the wild-type mice and a set of primers that flank the LoxP-common forward primer 5′-GAGATGGCGCAACGCAATTAAT-3′ and gene specific reverse primer 5′-ACACACTCCACTGAACAAAGTCAAGC-3′, to give a product of 437bp in homozygous mutants, with both products present in heterozygous littermates.

### Auditory Brainstem Response

To investigate auditory function and phenotype, ABR tests were performed on P20, P30, P40, and P60 mice using tone-burst stimuli. Briefly, mice were anaesthetized by intraperitoneal injection of xylazine (20 mg/ml^1^ at 5% v/v) and ketamine (100 mg/ml^−1^ at 10% v/v) administered at the rate of 0.1 ml per 10 g body mass, and placed in an acoustic chamber (MAC-1, Industrial Acoustic Company), as previously described [41].

### Scanning electron microscopy

The inner ears were dissected in cold PBS buffer shortly after mice were euthanized by C0_2_ inhalation. The temporal bone was removed prior to overnight fixation in glutaraldehyde (2.5 % v/v in PBS) at 4°C. The samples were alternately incubated in osmium tetroxide and thiocarbohydrazide after exposing the organ of Corti, as previously described [42]. After treatment, the samples were vacuum dried and mounted on a metal plate. Subsequently the samples were gold-coated at the Faculty of Life Sciences Electron Microscopy Unit at Tel Aviv University and imaged with a JSM 540A scanning electron microscope (Jeol).

### Western blot analysis

Cochlea and Huh7 cell protein lysates were prepared using Nonidet P-40 lysis buffer [150 mM NaCl,1.0% Nonidet P-40, Tris-Cl (50mM pH 8.0) protease inhibitor mixture, for 30 min on ice. The lysate was cleared by centrifugation at 13200 rpm for 15 min at 4 °C, and supernatant was recovered. Protein concentration was determined using the BCA protein determination reagent (Sigma), and 50 μg were resolved on an SDS/PAGE denaturing gel and transferred to a nitrocellulose membrane. Immunoblots were performed using the appropriate antibodies, and the membranes were developed using the Quantum ECL detection kit (K-12042-D20; Advansta). The immunoblot bands were quantified using ImageJ software, and the variation in protein loading was corrected by normalization to the levels of the indicated loading control protein such as tubulin. For IP, the primary antibody was incubated with protein A/G agarose beads (Santa Cruz Biotechnology, Dallas, TX, USA) at 4°C with mild shaking. 2 mg of cleared lysate was precleared with protein A/G agarose beads for 1 hour at 4°C and incubated overnight with antibody-conjugated protein A/G agarose beads at 4°C. Beads were recovered and washed 5 times with lysis buffer before resolving in SDS-PAGE. Subsequently IP was confirmed with the appropriate antibody.

### Cochlea protein extraction

Total protein from cochlea was extracted as previously described [43]. Briefly, 12 cochleas from wild-type P0 mice were dissected and lysed with 10% NP-40 protease inhibitor mixture, kept for 30 min on ice, and centrifuged at 13200 rpm for 15 min at 4°C, to harvest the supernatant. Protein concentration was determined using the BCA protein determination reagent (Sigma), and 60 μg were resolved on an SDS/PAGE gel and transferred to a nitrocellulose membrane. Immunoblots were performed using the appropriate antibodies. The membranes were developed using the WesternBright Quantam kit (K-12042-D20; Advansta, San Jose, CA, USA).

### Tail protein extraction

To confirm the genotyping, total protein was homogenized from the tails using BioVortexer (BioSpec Products, Bartlesville, OK, USA) and 120 μg of protein was resolved on an SDS/PAGE, as subjected to western blot analysis.

### Immunolocalization

Whole mount immunohistochemistry of inner ear was performed as previously described [44]. Briefly, the inner ears were dissected in cold PBS buffer shortly after P30 mice were euthanized by C0_2_ inhalation. Temporal bone was removed prior to overnight fixation in paraformaldehyde (4 % v/v in PBS) at 4 °C. The sensory epithelium was fine dissected, blocked, and permeabilized by incubation in blocking buffer (normal goat serum in 0.1% triton) for 2 hours at temperature. Samples were incubated with indicated primary antibody overnight at 4°C. After a brief wash in PBSX1, samples were incubated with secondary antibody for 2 hours at room temperature. To visualize F actin, this was followed by incubation for 1 hour at room temperature in phalloidin conjugated to Alexa Fluorophores (Life Technologies). The stained samples were mounted on Histobond microscope slides (Marienfeld GmbH) using Prolong Gold (Thermo Scientific) and dried overnight at room temperature. Image acquisition was performed with a confocal laser microscopy system (LSM800 Carl Zeiss).

### Endocochlear potential recording

Mice were anaesthetized by intraperitoneal injection of 50 mg/kg pentobarbital. The skin covering the neck was cut to expose the trachea. A tracheostomy was performed in order to maintain sufficient ventilation. The cochlea was exposed by a ventral approach and the tympanic bulla was gently picked to expose the cochlea. A drill was then used to expose the spiral ligament beneath the lateral wall. A glass pipette filled with 150 mM KCl was gradually inserted into the scala media through the spiral ligament while continuously recording the DC potential. The EP was defined as the delta between the potential recorded in the scala media and the one recorded on the spiral ligament. Potentials were amplified by OC-725C (Warner Instruments, CT, USA), digitized at 1 kHz using MiniDigi 1A (Molecular Devices, CA, USA) and analyzed using pCLAMP 9 (Molecular Devices).

### Antibodies

The following antibodies were used for this study: rabbit anti-striatin (IB: 1:1,000; IHC 1:250; Proteintech) mouse anti-striatin (IB: 1:1000; IHC: 1;250; BD Transduction Laboratories), mouse anti-Ctbp2 (IHC 1:250; BD Transduction Laboratories), rabbit anti-Myosin-VIIA (IHC 1:250; Proteus Biosciences), mouse anti-ZO1 (IHC: 1:100; Thermo Scientific), mouse anti-PP2A (IB:1:1000; Upstate Biotechnology), rabbit anti-striatin 4 (IHC: 1:250; Abcam), rat anti-Ecad (IHC:1:250; Santa Cruz), Phallodin-488 (IHC: 1:1000; Abcam), mouse anti-Alexa fluor 594 (IHC: 1:250; Abcam), rabbit anti-Alexa fluor 633 (IHC: 1:250; Invitrogen); mouse anti-tubulin (IB 1:10,000; Sigma) was used as a loading control.

### Quantification of ribbon synapses

Whole mount immunohistochemistry was performed as described. The region between the first and second turn of the cochlea was dissected and stained with Ctbp2 and myosin VIIA antibody. The samples were carefully mounted to avoid excessive pressure from the coverslip that can squeeze the tissue. Image acquisition was performed with a confocal laser microscopy system (LSM800, Carl Zeiss, Oberkochen, Germany). The Z stacks of the images were exported to Imaris software (Zurich, Switzerland). All the analyses were performed using the same settings in Imaris. For P17, 4-11 IHCs, and for P35, 5-10 IHCs, from each cochlea were quantified after exporting the file to Imaris.

## Acknowledgements

The research was supported by a US-Israel Binational Science Foundation grant 2017173 (RRA), National Institutes of Health/NIDCD R01DC011835 (KBA), and a Cincinnati Children’s Hospital Medical Center – Tel Aviv University Sackler Faculty of Medicine Grant for Collaborative Research (RL, RRA). We thank Ana Turchetti-Maia and Kevin Ohlemiller for their advice for EP recordings, Vered Holdengreber for help with electron microscopy, and Ronen Siman-Tov for help with the mice.

## Author contributions

KBA and RRA conceived the project. RAL generated the Striatin knockout mice. PTNP designed and performed the genotyping, dissections of inner ears, ABR, SEM, confocal imaging, Western blots, and quantification of ribbon synapses. MR and ST performed the endocochlear potential experiments. ST and TK-B aided with dissections and SEM imaging analysis. MC and AAD aided with image analysis, experimental design and organizing the data. PTNP, ST, KBA, and RRA wrote the manuscript. KBA and RRA supervised the work.

## Conflict of interest

The authors declare that they have no competing interest.

## Supporting Information

### Expanded View Figures

**Figure EV1.**
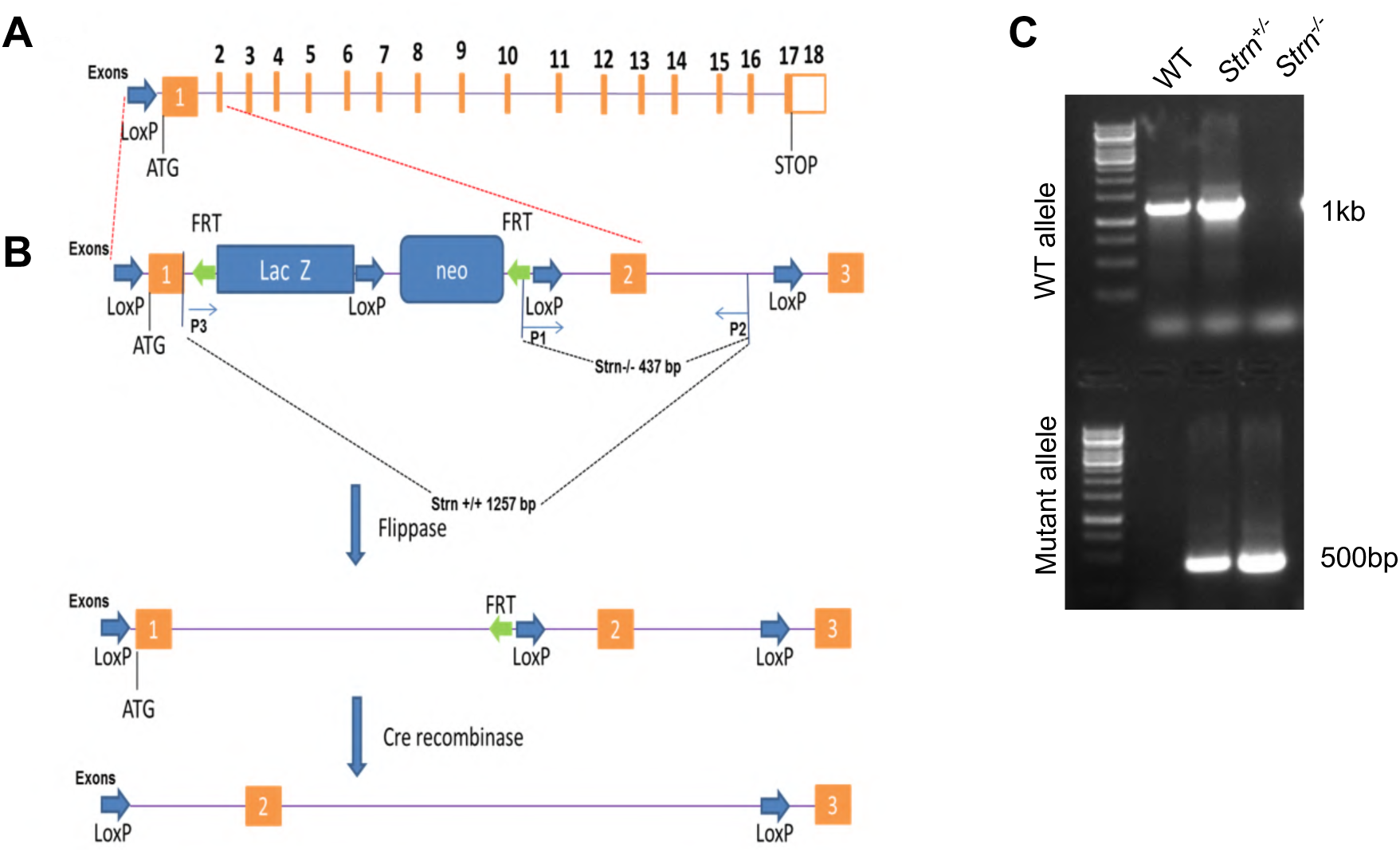
Generation of the striatin knockout mice line. A Schematic representation of the striatin gene. Rectangles represent exons, with coding sequence colored in orange. B The first and second exons are shown. Green arrowheads are FRT sites. Blue arrowheads denote LoxP sites. Recombination of the FRT sites flanking the Neo results in the floxed Strn allele. Recombination of the LoxP sites removes the ATG start site in exon 1 resulting in the *Strn* null allele P1, P2 and P3 represent the location of the primers used for *Strn* genotyping. C For genotyping, tails were excised from mice and genomic DNA was extracted and subjected to PCR analysis. Primers were designed to generate amplicons of 1.2kb for the WT and 450bp for the null mutant.

**Figure EV2.**
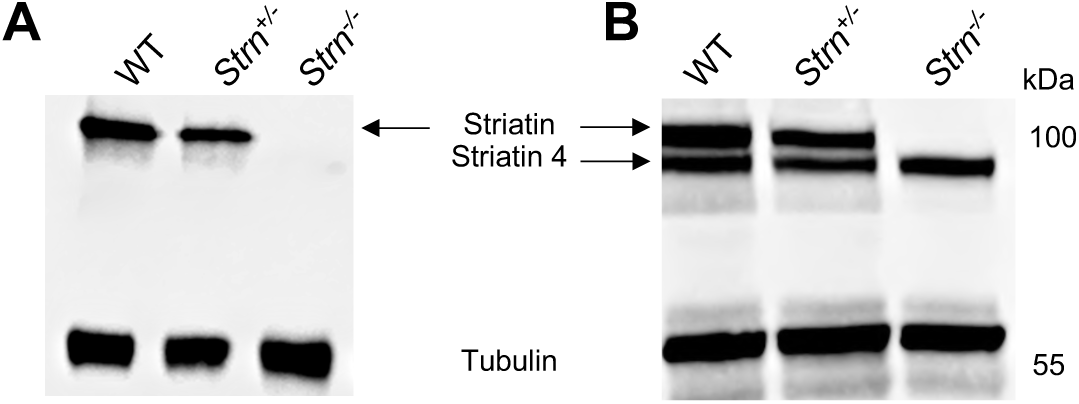
The expression of Striatin 4 is unaffected in *Strn*^*-/-*^ mice. Total protein extracted from tails was resolved and immunoblotted for western blot analysis, using the indicated antibodies. A Validation of mice genotyping using an anti-striatin antibody that detects only the Strn1 isoform. B Western blot analysis shows that the expression of striatin 4 is unaffected in *Strn*^*-*^ */*^*-*^ mutants at P58. Tubulin was used as loading control.

**Figure EV3.**
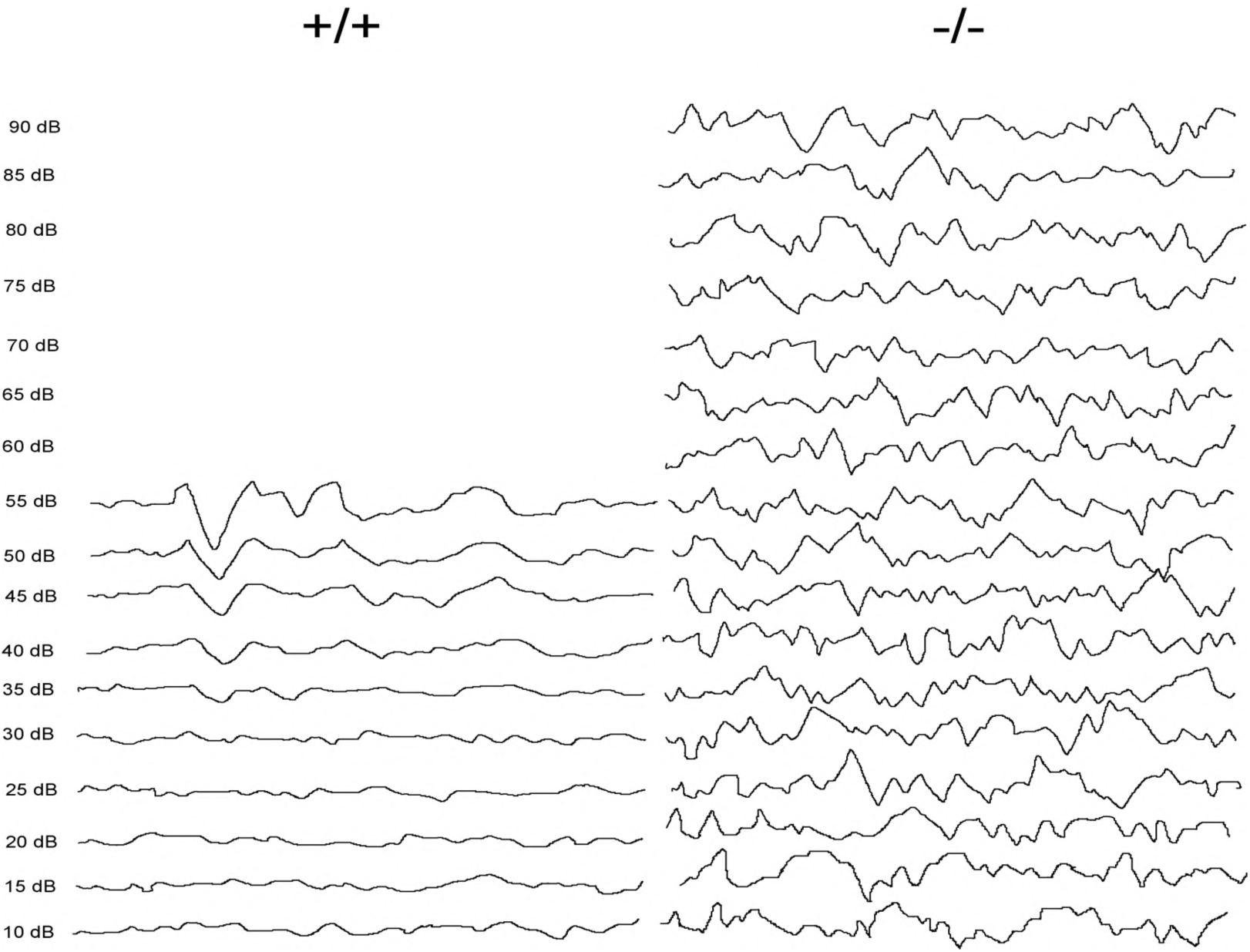
Representative ABR waves at 30 KHz showing increased threshold for *Strn*^*-/-*^ as compared to the control.

## References

1. Hwang J, Pallas DC (2014) STRIPAK complexes: structure, biological function, and involvement in human diseases. Int J Biochem Cell Biol 47: 118–148

2. Kuck U, Radchenko D, Teichert I (2019) STRIPAK, a highly conserved signaling complex, controls multiple eukaryotic cellular and developmental processes and is linked with human diseases. Biol Chem pii: /j/bchm.ahead-of-print/hsz-2019-0173/hsz-2019-0173.xml.

3. Shi Z, Jiao S, Zhou Z (2016) STRIPAK complexes in cell signaling and cancer. Oncogene 35: 4549–4557

4. Castets F, Rakitina T, Gaillard S, Moqrich A, Mattei MG, Monneron A (2000) Zinedin, SG2NA, and striatin are calmodulin-binding, WD repeat proteins principally expressed in the brain. J Biol Chem 275: 19970–19977

5. Moreno CS, Park S, Nelson K, Ashby D, Hubalek F, Lane WS, Pallas DC (2000) WD40 repeat proteins striatin and S/G(2) nuclear autoantigen are members of a novel family of calmodulin-binding proteins that associate with protein phosphatase 2A. J Biol Chem 275: 5257–5263

6. Lahav-Ariel L, Caspi M, Nadar-Ponniah PT, Zelikson N, Hofmann I, Hanson KK, Franke WW, Sklan EH, Avraham KB, Rosin-Arbesfeld R (2019) Striatin is a novel modulator of cell adhesion. FASEB J 33: 4729–4740

7. Dallos P (1992) The active cochlea. J Neurosci 12: 4575–4585

8. Kalluri R, Monges-Hernandez M (2017) Spatial Gradients in the Size of Inner Hair Cell Ribbons Emerge Before the Onset of Hearing in Rats. J Assoc Res Otolaryngol 18: 399–413

9. Liberman LD, Wang H, Liberman MC (2011) Opposing gradients of ribbon size and AMPA receptor expression underlie sensitivity differences among cochlear-nerve/hair-cell synapses. J Neurosci 31: 801–808

10. Liberman MC (1982) Single-neuron labeling in the cat auditory nerve. Science 216: 1239–1241

11. Yin Y, Liberman LD, Maison SF, Liberman MC (2014) Olivocochlear innervation maintains the normal modiolar-pillar and habenular-cuticular gradients in cochlear synaptic morphology. J Assoc Res Otolaryngol 15: 571–583

12. Hickman TT, Liberman MC, Jacob MH (2015) Adenomatous Polyposis Coli Protein Deletion in Efferent Olivocochlear Neurons Perturbs Afferent Synaptic Maturation and Reduces the Dynamic Range of Hearing. J Neurosci 35: 9236–9245

13. Frost A, Elgort MG, et al (2012) Functional repurposing revealed by comparing S. pombe and S. cerevisiae genetic interactions. Cell 149: 1339–1352

14. Jain BP, Pandey S, Saleem N, Tanti GK, Mishra S, Goswami SK (2017) SG2NA is a regulator of endoplasmic reticulum (ER) homeostasis as its depletion leads to ER stress. Cell Stress Chaperones 22: 853–866

15. Tanti GK, Goswami SK (2014) SG2NA recruits DJ-1 and Akt into the mitochondria and membrane to protect cells from oxidative damage. Free Radic Biol Med 75: 1–13

16. Ranum PT, Goodwin AT, Yoshimura H, Kolbe DL, Walls WD, Koh JY, He DZZ, Smith RJH (2019) Insights into the Biology of Hearing and Deafness Revealed by Single-Cell RNA Sequencing. Cell Rep 26: 3160–3171 e3163

17. Chessum L, Matern MS, et al (2018) Helios is a key transcriptional regulator of outer hair cell maturation. Nature 563: 696–700

18. Rudnicki A, Isakov O, Ushakov K, Shivatzki S, Weiss I, Friedman LM, Shomron N, Avraham KB (2014) Next-generation sequencing of small RNAs from inner ear sensory epithelium identifies microRNAs and defines regulatory pathways. BMC Genomics 15: 484

19. Kamitani T, Sakaguchi H, et al (2015) Deletion of Tricellulin Causes Progressive Hearing Loss Associated with Degeneration of Cochlear Hair Cells. Sci Rep 5: 18402

20. Kitajiri S, Katsuno T, Sasaki H, Ito J, Furuse M, Tsukita S (2014) Deafness in occludin-deficient mice with dislocation of tricellulin and progressive apoptosis of the hair cells. Biol Open 3: 759–766

21. Higashi T, Katsuno T, Kitajiri S, Furuse M (2015) Deficiency of angulin-2/ILDR1, a tricellular tight junction-associated membrane protein, causes deafness with cochlear hair cell degeneration in mice. PLoS One 10: e0120674

22. Kazmierczak M, Harris SL, Kazmierczak P, Shah P, Starovoytov V, Ohlemiller KK, Schwander M (2015) Progressive Hearing Loss in Mice Carrying a Mutation in Usp53. J Neurosci 35: 15582–15598

23. Kazmierczak-Baranska J, Peczek L, Przygodzka P, Cieslak MJ (2015) Downregulation of striatin leads to hyperphosphorylation of MAP2, induces depolymerization of microtubules and inhibits proliferation of HEK293T cells. FEBS Lett 589: 222–230

24. Patuzzi R (2011) Ion flow in stria vascularis and the production and regulation of cochlear endolymph and the endolymphatic potential. Hear Res 277: 4–19

25. Quraishi IH, Raphael RM (2008) Generation of the endocochlear potential: a biophysical model. Biophys J 94: L64–66

26. Xiong H, Chu H, et al (2011) Conservation of endocochlear potential in mice with profound hearing loss induced by co-administration of kanamycin and furosemide. Lab Anim 45: 95–102

27. Florian P, Amasheh S, Lessidrensky M, Todt I, Bloedow A, Ernst A, Fromm M, Gitter AH (2003) Claudins in the tight junctions of stria vascularis marginal cells. Biochem Biophys Res Commun 304: 5–10

28. Kitajiri S, Miyamoto T, et al (2004) Compartmentalization established by claudin-11-based tight junctions in stria vascularis is required for hearing through generation of endocochlear potential. J Cell Sci 117: 5087–5096

29. Viquez NM, Fuger P, Valakh V, Daniels RW, Rasse TM, DiAntonio A (2009) PP2A and GSK-3beta act antagonistically to regulate active zone development. J Neurosci 29: 11484–11494

30. Khimich D, Nouvian R, Pujol R, Tom Dieck S, Egner A, Gundelfinger ED, Moser T (2005) Hair cell synaptic ribbons are essential for synchronous auditory signalling. Nature 434: 889–894

31. Pisciottano F, Cinalli AR, Stopiello JM, Castagna VC, Elgoyhen AB, Rubinstein M, Gomez-Casati ME, Franchini LF (2019) Inner Ear Genes Underwent Positive Selection and Adaptation in the Mammalian Lineage. Mol Biol Evol 36: 1653–1670

32. Breitman M, Zilberberg A, Caspi M, Rosin-Arbesfeld R (2008) The armadillo repeat domain of the APC tumor suppressor protein interacts with Striatin family members. Biochim Biophys Acta 1783: 1792–1802

33. Tran H, Hamada F, Schwarz-Romond T, Bienz M (2008) Trabid, a new positive regulator of Wnt-induced transcription with preference for binding and cleaving K63-linked ubiquitin chains. Genes Dev 22: 528–542

34. Neisch AL, Neufeld TP, Hays TS (2017) A STRIPAK complex mediates axonal transport of autophagosomes and dense core vesicles through PP2A regulation. J Cell Biol 216: 441–461

35. Niceta M, Stellacci E, et al (2015) Mutations Impairing GSK3-Mediated MAF Phosphorylation Cause Cataract, Deafness, Intellectual Disability, Seizures, and a Down Syndrome-like Facies. Am J Hum Genet 96: 816–825

36. Xie WR, Jen HI, Seymour ML, Yeh SY, Pereira FA, Groves AK, Klisch TJ, Zoghbi HY (2017) An Atoh1-S193A Phospho-Mutant Allele Causes Hearing Deficits and Motor Impairment. J Neurosci 37: 8583–8594

37. Zhai X, Liu C, Zhao B, Wang Y, Xu Z (2018) Inactivation of Cyclin-Dependent Kinase 5 in Hair Cells Causes Hearing Loss in Mice. Front Mol Neurosci 11: 461

38. Goudreault M, D’Ambrosio LM, et al (2009) A PP2A phosphatase high density interaction network identifies a novel striatin-interacting phosphatase and kinase complex linked to the cerebral cavernous malformation 3 (CCM3) protein. Mol Cell Proteomics 8: 157–171

39. Kean MJ, Ceccarelli DF, et al (2011) Structure-function analysis of core STRIPAK Proteins: a signaling complex implicated in Golgi polarization. J Biol Chem 286: 25065–25075

40. Scheffer DI, Shen J, Corey DP, Chen ZY (2015) Gene Expression by Mouse Inner Ear Hair Cells during Development. J Neurosci 35: 6366–6380

41. Horn HF, Brownstein Z, et al (2013) The LINC complex is essential for hearing. J Clin Invest 123: 740–750

42. Hunter-Duvar IM (1978) A technique for preparation of cochlear specimens for assessment with the scanning electron microscope. Acta Otolaryngol Suppl 351: 3–23

43. Bhonker Y, Abu-Rayyan A, et al (2016) The GPSM2/LGN GoLoco motifs are essential for hearing. Mamm Genome 27: 29–46

44. Dror AA, Politi Y, et al (2010) Calcium oxalate stone formation in the inner ear as a result of an Slc26a4 mutation. J Biol Chem 285: 21724–21735

